# A comparative study between state-of-the-art MRI deidentification and AnonyMI, a new method combining re-identification risk reduction and geometrical preservation

**DOI:** 10.1101/2021.07.30.454335

**Authors:** Ezequiel Mikulan, Simone Russo, Flavia Maria Zauli, d’Orio Piergiorgio, Sara Parmigiani, Jacopo Favaro, William Knight, Silvia Squarza, Pierluigi Perri, Francesco Cardinale, Pietro Avanzini, Andrea Pigorini

## Abstract

Deidentifying MRIs constitutes an imperative challenge, as it aims at precluding the possibility of re-identification of a research subject or patient, but at the same time it should preserve as much geometrical information as possible, in order to maximize data reusability and to facilitate interoperability. Although several deidentification methods exist, no comprehensive and comparative evaluation of deidentification performance has been carried out across them. Moreover, the possible ways these methods can compromise subsequent analysis has not been exhaustively tested. To tackle these issues, we developed AnonyMI, a novel MRI deidentification method, implemented as a user-friendly 3D Slicer plugin-in, which aims at providing a balance between identity protection and geometrical preservation. To test these features, we performed two series of analyses on which we compared AnonyMI to other two state-of-the-art methods, to evaluate, at the same time, how efficient they are at deidentifying MRIs and how much they affect subsequent analyses, with particular emphasis on source localization procedures. Our results show that all three methods significantly reduce the re-identification risk but AnonyMI provides the best geometrical conservation. Notably, it also offers several technical advantages such as a user-friendly interface, multiple input-output capabilities, the possibility of being tailored to specific needs, batch processing and efficient visualization for quality assurance.

## 1. Introduction

A growing amount of evidence points to open science as the most efficient strategy for understanding the structural and functional organization of the human brain (Milham et al., 2018). This notion has driven several recent unprecedented large-scale collaborative endeavors in neuroscience (Amunts et al., 2019; Mott et al., 2018). As a natural consequence, in the last years, data sharing has promptly become a fundamental pillar of contemporary neuroscience. Indeed, it allows otherwise impossible large-scale and multi-centric analyses and also promotes transparency and reproducibility (Ascoli et al., 2017). Data sharing also maximizes the scientific profit that can be obtained from the data (Brakewood and Poldrack, 2013), reduces the economic cost of new studies, and provides valuable resources for researchers without access to expensive data acquisition equipment (Poline et al., 2012).

However, human data sharing also entails some risks, mostly related to privacy and confidentiality of personal information and metadata. Indeed, protecting the privacy and confidentiality of research subjects is not always granted, even though considerable efforts have been directed to establish data deidentification protocols and tools, and data protection legislation (Kalkman et al., 2019). Researchers must ensure that they adhere to the applicable data protection legislation. Within Europe, the General Data Protection Regulation (GDPR; EU 679/2016) establishes the law on data protection issues. Brought into force in May 2018, the GDPR covers only those data where there is the possibility of identifying the data subject from its contents – and it applies a number of restrictions, such as how data can be shared and stored, who can access them, and how it might be ultimately used and reused. If it is not possible to re-identify subjects from the data, then that data falls outside of the provisions of GDPR – and so such restrictions do not apply. Thus, there is a pressing need, then, to classify which data entails (or not) the risk of identification of the participant.

The problem of possible re-identification of participants is fundamental for data types containing explicit physical features, as in the case of Magnetic Resonance Imaging (MRI). Typical MRI acquisitions in neuroscience, aside from the brain, contain detailed representations of the facial features of the participants (Figure 1) that can be used to create accurate three-dimensional representations of the subject’s faces (Prior et al., 2009). To date, several solutions exist, which range from extracting the brain and sharing only this part of the volumes (Kalavathi and Prasath, 2016), to facial masking and facial removal (Bischoff-Grethe et al., 2007), but each of these methods has its downsides. For example, different masking and defacing procedures can affect subsequent steps of analysis and even lead to failure (de Sitter et al., 2020). Moreover, when MRIs are shared as part of EEG studies, brain extraction may impede accurate source modelling analyses that take advantage of head geometries (Hallez et al., 2007), and current facial masking or removal techniques may induce geometrical distortions.

**Figure 1.**
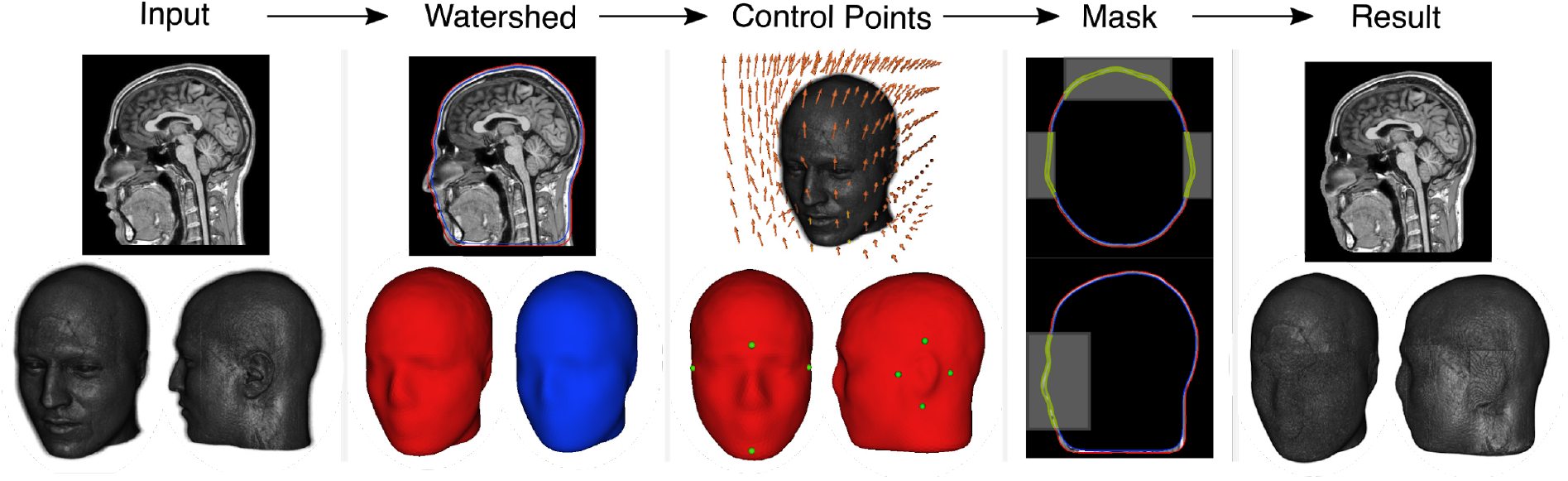
AnonyMI. Illustration of the deidentification procedure. The deidentification procedure is performed by first using a watershed algorithm to obtain 3D reconstructions of the skin and skull of the subject, then aligning the subject’s MRI to a template that contains control points for the face and ears (or indicating them manually), and finally applying a mask to the intersection of these control points and the skin and skull surfaces. The MRI shown in this example is from a subject that provided informed consent for it to be shown.

Few studies have quantitatively tested the re-identification risk and the geometrical preservation offered by these methods (Budin et al., 2008; de Sitter et al., 2020; Mazura et al., 2012; Prior et al., 2009), and no comprehensive comparison among them has been performed. Treating these two aspects simultaneously is of paramount importance not only for investigators and research participants but also for properly informed legislation regarding data protection. Indeed, without such information, it is impossible to know whether currently accepted methods comply with the GDPR requirements and, at the same time, to what extent the geometrical properties of the subjects can be preserved without risking re-identification.

In that vein, here we present a new method for deidentifying MRIs - called AnonyMI- and we compare it with two state-of-the-art deidentification methods by evaluating, for the first time, their performance with tailored behavioral tasks. With respect to other methods, AnonyMI aims at preserving most of the geometrical properties of the subject’s head whilst effectively deidentifying the MRI. It takes advantage of the Boundary Element Method (BEM; Hallez et al., 2007), which is a standard procedure for creating head models for source localization of scalp EEG activity. Specifically, AnonyMI employs the same three-dimensional surfaces that are used for source modelling (Akalin-Acar and Gençer, 2004; Hamalainen and Sarvas, 1987) to mask the subjects’ facial features, which results in the same surfaces being obtained when re-computing the surfaces on the anonymized MRIs.

To test the performance of AnonyMI, we performed 2 sets of analyses, in both cases comparing it with other two state-of-the-art methods: PyDeface and Maskface (Gulban et al., 2019; Milchenko and Marcus, 2013). Specifically, first we performed a behavioral validation in which we tested and compared, on two experiments, how effectively each method deidentified the MRIs by asking participants to identify subjects from 3D reconstructions of their MRIs. Second, we performed a geometrical evaluation in which we measured how much each method introduces spatial distortions and affects subsequent processing stages, and also compared their performance when being employed for source localization analysis using ground-truth data (Mikulan et al., 2020). Our results show that AnonyMI outperforms the other two methods in terms of geometrical conservation. Most importantly, we also show that AnonyMI, similarly to the other two tested deidentification methods, allows taking the re-anonymization risk close to chance level.

## 2. Methods

### 2. 1. AnonyMI

AnonyMI is an MRI deidentification tool that uses 3D surface modelling in order to deidentify MRIs whilst retaining as much geometrical information as possible. It can be run automatically or manually, which allows precise tailoring for specific needs. It is distributed as a plug-in of 3D Slicer, a widely used, open-source, stable, and reliable image processing software (Fedorov et al., 2012). It leverages the power of this platform for reading and saving images which makes it applicable on almost any MRI filetype, including all the most commonly used formats (e.g. DICOM, Nifti, Analyze, etc.). As it uses an algorithm from Freesurfer (Dale et al., 1999) it currently runs on Linux and Mac platforms, but native Windows support will be possible in the near future and is indeed possible at the present with an elaborate setup. This algorithm is provided inside the toolbox and therefore does not require installing Freesurfer. Importantly, AnonyMI operates on the image data and does not apply changes to the file header, which might contain sensitive information. Several options exist for removing sensitive data from MRI file headers (Kushida et al., 2012). However, AnonyMI, by default, saves images in NIfTI format which is less prone to contain private information. Nevertheless, it is the responsibility of the user to ensure that the file header is free of sensitive data.

The procedure involves three main steps (Figure 1). First, the *watershed* algorithm, taken from Freesurfer (Dale et al., 1999; Ségonne et al., 2004), is applied in order to obtain 3D reconstructions of the skin and skull of the subject. It is worth mentioning that these are the same surfaces that are often used in the creation of forward solutions for source modelling (Hallez et al., 2007). Next, the location of the face and ears are determined by performing a non-linear registration between the subject’s MRI and a template, taken from the IXI dataset (Avants and Tustison, 2018), on which these control points have already been marked. The location of these points can also be done manually by taking advantage of the extensive rendering and interactive capabilities of 3D Slicer. This allows extending the areas to be anonymized at will, in order to cover specific parts that might increase the subject’s identification risk (e.g. scars, malformations, etc.). In addition, creating a new template is fast and straightforward and can be done by simply importing the image to be used and marking the face and ears control points. This permits the creation of population-specific templates (e.g. age-specific) that can then be used to perform automatic anonymization. Finally, a subject-specific mask is created by taking the intersection of the control points and the skin and skull surfaces, which is filled with random numbers that follow the distribution of voxel intensities inside the space between the two surfaces. The resulting volume is an anonymized MRI that preserves most of the geometrical information of the subject.

### 2.2. Validation

In order to assess the performance of our method and validate its results, we performed two series of analyses. First, we assessed the deidentification level achieved in two behavioral experiments. Second, we evaluated the similarity of the deidentified MRIs with respect to the original versions and tested how using deidentified MRIs affects EEG source localization results. In all cases, we compared the results obtained with our method to those obtained with other two state-of-the art methods, PyDeface and Maskface. Data analyses were performed in Python and statistical analyses were performed in R (R Core Team, 2019).

#### 2.2.1. Behavioral Validation

We conducted two behavioral experiments to evaluate how much different deidentification methods reduce the probability of identification of a subject from a 3D reconstruction of his/her MRI. On each trial of the first experiment, participants were presented with one photograph and then four deidentified MRIs, and they had to indicate to which of them the photograph belonged. In the second experiment, the procedure was inverted, that is, participants were presented with four photographs and then one deidentified MRI, and had to indicate to which photograph the MRI belonged. The two versions of the experiment aimed to provide ecologically valid estimates of the deidentification performance of each method as they correspond to trying to identify a person from a series of available MRIs, and vice versa, trying to identify an MRI from a series of available candidates.

##### 2.2.1.1. Experiment 1

###### Stimuli

Stimuli consisted in 3D reconstructions of MRIs from 15 subjects (F=7, 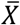 age = 33.8, *sd* age = 7.28) that were deidentified with three different methods (PyDeface, Maskface and AnonyMI) and their corresponding photographs (front, left and right views). Reconstructions were obtained from 3D Slicer’s Volume Rendering module and photographs were acquired with a light grey background. All images were post-processed using GIMP, where they were converted to grayscale, centered, aligned (horizontally across the eyeline), cropped to the same size, and finally normalized using the histogram equalization technique (see Supplementary Figure 1.a). In the case of MRI reconstructions that presented noise around the edges of the head, they were manually cleaned by background cloning (see Supplementary Figure 1.b).

###### Participants & Procedure

A total of 31 subjects participated in the experiment (F = 19 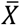 age = 20.16, *sd* age = 0.58). The experiment was performed in three consecutive groups in the same room with one desktop computer per participant. Participants were instructed that on each trial they would first see the photograph of a person, followed by four 3D reconstructions of deidentified MRIs, and that their task was to indicate, using the numbers from 1 to 4 on the keyboard, which was the MRI that corresponded to the photograph they had seen. Each participant performed a total of 135 trials in ∼25 minutes. The experiment was programmed using PsychoPy3 (Peirce et al., 2019). At the beginning of the experiment, the software created a random sequence of trials for each participant by choosing for each trial a photograph, its corresponding MRI and 3 random sex-matched distractor MRIs. Each trial started with a fixation cross (1 second), followed by the presentation of the photograph at the center of the screen (3 seconds), followed by 4 MRI reconstructions displayed on a 2 by 2 grid, each one with a number from 1 to 4 on top (Figure 2.a). Participants could take all the time they needed on each trial to respond.

**Figure 2.**
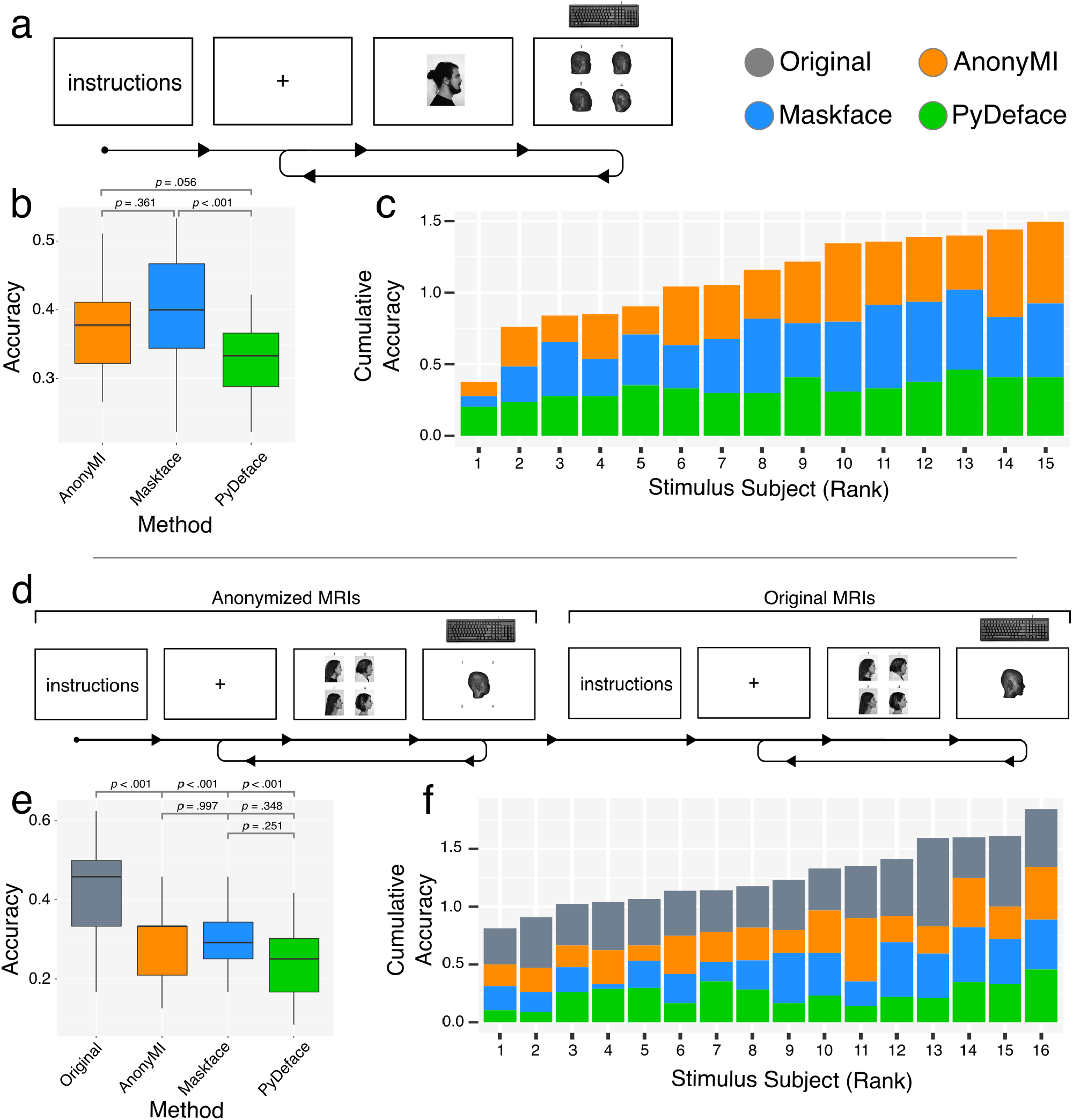
Behavioral results. a) Illustration of the experimental procedure of Experiment 1. b) Boxplot of mean accuracy by participant for each deidentification method of Experiment 1. c) Cumulative accuracy by stimulus subject for each deidentification method of Experiment 1. d) Illustration of the experimental procedure of Experiment 2. e) Boxplot of mean accuracy by participant for each deidentification method of Experiment 2. f) Cumulative accuracy by stimulus subject for the original MRIs and for each deidentification method of Experiment 2. Statistical analysis of panel b and e were carried out employing binomial generalized linear mixed effects models with random intercept per participant. Post-hoc comparisons were performed using estimated marginal means and Tukey *p*-value adjustment For multiple comparisons.

###### Analysis

In order to quantify the differences in the deidentification performance of these methods, we performed a mixed-effects logistic regression analysis with participant as random factor, as it is one of the recommended approaches for accuracy data (Dixon, 2008; Jaeger, 2008). This analysis was performed using the *lme4* package (Bates et al., 2015), which provides p-values via Wald tests. We defined *deface* as the reference level (as it is arguably the most performant method due to the complete removal of facial attributes) and assessed how much the probability of correct responses changed by using the other two methods. Model fits were judged by visual inspection of simulated residuals plot (i.e. diagonal patterns in Q-Q plots) using the DHARMa package with 10^3^ replications. Post-hoc comparisons were performed using estimated marginal means and Tukey p-value adjustment for multiple comparisons.

##### 2.2.1.2. Experiment 2

###### Stimuli

The same stimuli of Experiment 1 were used, with the addition of one more MRI (and its corresponding photographs), adding up to a total of 16 subjects (F=7, 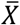 age = 34.1 *sd* age = 7.11), and 40 photographs from new subjects that were used as distractors (F=14, 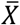 age = 38.6, *sd* age = 6.81). All images were post-processed as described in Experiment 1. Test and control stimuli matching was performed separately for each participant as each one was presented with only a subset of the stimuli pool (see below).

###### Participants & Procedure

A total of 24 subjects participated in the experiment (F = 11; 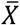 age = 30.5, *sd* age = 6.57). The experiment was programmed in PsychoPy and it was performed online by participants through the Pavlovia platform (https://pavlovia.org). They were instructed that on each trial they would first see four photographs from different persons followed by a 3D reconstruction of an MRI, and that their task was to indicate, using the numbers from 1 to 4 on the keyboard, to which of the subjects they had seen it belonged. The numbers that indicated each stimulus remained on the screen while the MRI image was being presented, in order to ease the memory load of the task. Participants completed a total of 96 trials in ∼20 minutes. One hundred random combinations of stimuli were created offline and each participant was assigned one of them using a random number generator at the beginning of the experiment. These combinations of stimuli were created by selecting 6 males and 6 females with MRIs, and 6 males and 6 females from the distractors pool. For each deidentification method, each of these stimuli subjects was presented the same number of times (once from the front view and once from one of the lateral views) in random combinations. After all trials using deidentified MRIs were completed, a control block was presented in which stimuli were created in the same way but using the 3D reconstructions of the original MRIs (i.e. non deidentified) in order to obtain a reference score for the performance of each participant. The control block was presented at the end of the experiment, instead of aside to the deidentification methods, in order to avoid that subjects recognized distinctive features on the non-anonymized stimuli that could have later served as cues in anonymized trials. On each trial subjects were presented with a fixation cross (1 second), followed by the four photographs (7 seconds) on a 2 by 2 grid, followed by the MRI (maximum of 10 seconds to respond; Figure 2.d).

###### Analysis

The same analyses as in Experiment 1 were used (see above), with the only difference of using the performance with the original MRIs as reference level. Namely, we assessed how much was the probability of correct responses reduced by using deidentified MRIs with respect to the performance with non-deidentified MRIs.

#### 2.2.2. MRI geometrical Similarity

In order to assess the geometrical similarity between the original and deidentified MRIs, we performed three analyses on a publicly available dataset (Souza et al., 2018), from which we randomly selected 75 MRIs that included data from 3 different scanner manufacturers (25 Siemens; 25 Phillips; 25 General Electric). First, we computed the Jaccard Similarity, which is computed as the ratio between the intersection and the union of two sets (Jaccard, 1912; Milchenko and Marcus, 2013). To this end, performed skull-stripping, using the ANTs toolbox (Avants et al., 2011), on the original and deidentified MRIs and calculated the similarity between each deidentified MRI and its corresponding original MRI. The skull-stripping procedure failed (i.e. removed brain tissue) on 8 of the original MRIs and therefore these were removed from the analysis. The template used for skull-stripping for AnonyMI and Maskface was the IXI template while the one used for PyDeface was the OASIS template from ANTs, as using a non-defaced template for the defaced images would have unfairly reduced its performance (Tustison et al., 2014). The comparison among methods was performed using pairwise Wilcoxon Rank Sum tests with Holm-Bonferroni correction for multiple comparisons.

Next, we employed the watershed algorithm from Freesurfer to create 3D reconstructions of the brain, inner skull, outer skull and outer skin, which are the same that are used for EEG/MEG source modelling. We assessed how many anonymized MRIs yielded usable reconstructions, that is, did not present deformations as appraised by visual inspection (Figure 3). We tested if the different deidentification methods produced significantly different numbers of usable reconstructions using Cochran’s Q test followed by post-hoc pairwise McNemar’s Chi-squared tests with Holm-Bonferroni correction for multiple comparisons.

**Figure 3.**
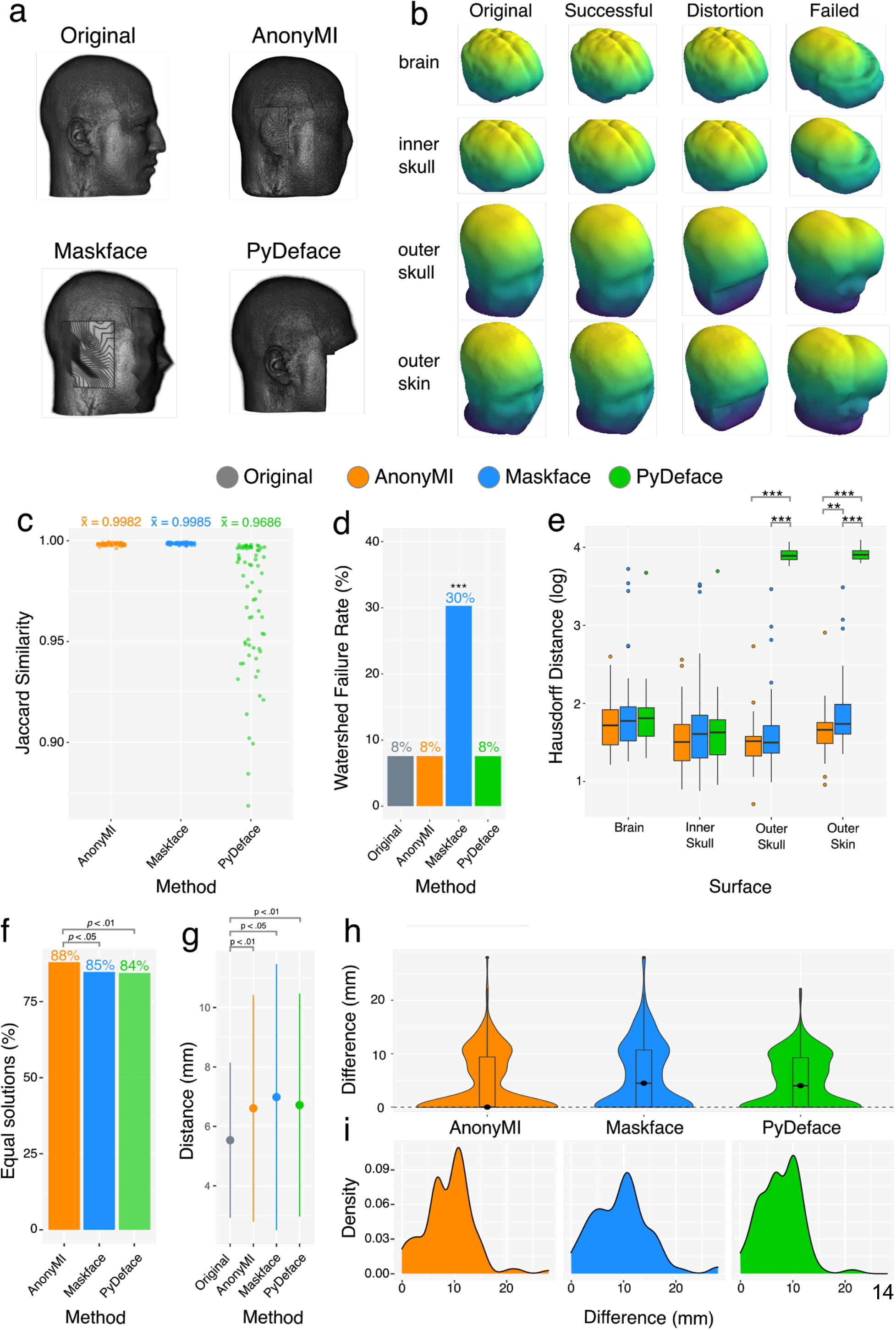
MRI similarity and source localization analyses. a) Example of 3D reconstructions of an original MRI and the three deidentification methods under evaluation. b) Example of original, successful, distorted and failed surface reconstructions obtained employing the watershed algorithm, c) Results of Jaccard Similarity between skull-stripped MRIs between the original image and each deidentification method. d) Percentage of failed watershed reconstructions (as depicted in rightmost column of *panel b*). Statistical analyses were performed using Cochran’s Q test and pairwise McNemar tests with Holm-Bonferroni correction for multiple comparisons. e) Results of the Hausdorff Distance analysis between the surfaces obtained with the original MRIs and those obtained with deidentified MRIs. Only images whose watershed reconstructions did not fail in any of the three deidentification methods were employed in this analysis. Statistical analyses were performed using pairwise Wilcoxon Rank Sum tests within surfaces with Holm-Bonferroni multiple comparisons correction. f) Percentage of solutions on which the reconstructed source was equal to the source obtained using the original MRI for each deidentification method. Statistical analyses were performed using Cochran’s Q test and pairwise McNemar tests with Holm-Bonferroni correction for multiple comparisons. g) Mean and standard deviation of distances to real source when using the original and deidentified MRIs. h) Violin plot and boxplot of difference between the source obtained with the original MRI and with each deidentification method when considering the solutions on which at least one method was different from the original solution. i) Density plot of difference with respect to the source obtained with the original MRI when considering only the different solutions for each method.

Finally, we computed the Hausdorff Distance (Huttenlocher et al., 1993), a measure of similarity of 3D models, between each of the 3D reconstructions of deidentified MRIs that did not present deformations and their corresponding reconstructions obtained from the original MRIs. We tested if these distances were different across deidentification methods for each of the four surfaces (brain, inner skull, outer skull and outer skin) using pairwise Wilcoxon Rank Sum tests within surfaces with Holm-Bonferroni multiple comparisons correction.

#### 2.2.3. Source Localization

In order to assess how much the geometrical distortions induced by the deidentification procedures affect source localization results we used the *LocalizeMI* dataset (Mikulan et al., 2020), which contains ground-truth data for assessing source localization performance, and tested how much the localization accuracy varied as a function of the deidentification method with respect to the localization obtained using the original MRI.

We used data from simultaneous HD-EEG (High-Density Electroencephalography) recordings and intracranial stimulation in order to assess how much results varied by employing deidentified MRIs in the creation of forward models. The description of the experimental procedure, pre-processing and source localization procedures can be found in the open-access publication that accompanies the dataset (Mikulan et al., 2020) and the corresponding scripts are also publicly available. Here, we repeated the analysis with all the parameter’s configurations that yielded optimal solutions for each of the 61 stimulation sessions, but this time using forward models created with deidentified MRIs. We then computed the percentage of solutions that resulted in the same localization error as the one obtained with the original MRIs. We tested if the different deidentification methods produced significantly different percentages of equal solutions using Cochran’s Q test followed by post-hoc pairwise McNemar’s Chi-squared tests with Holm-Bonferroni correction for multiple comparisons. Next, on all the configurations that did not result in the same localization error as in the original case for at least one method, we calculated the distance from the real source and tested how different it was from the distance obtained with the original MRIs by means of a mixed effects model. Specifically we used the log-transformed distance to the real source as dependent variable, each MRI deidentification method as predictor with the original MRIs as reference level, and subject as random factor (intercept and slope).

## 2. Results

### 3.1. Behavioral Validation

#### 3.3.1. Experiment 1

The overall performance (i.e. percentage of correct trials) was 33% for PyDeface, 37% for AnonyMI and 40% for Maskface. The mixed effects logistic regression analysis showed that AnonyMI increased the probability of correct responses (β = 0.18, *p* < .05) with respect to PyDeface, but this increase was smaller than in the case of Maskface (β = 0.28, *p* < .001). See Supplementary Table 1 for a complete report of this analysis. Interestingly the overall performance across stimulus subjects ranged from 12% to 49% (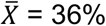, *sd* = 10%), showing that there were subjects that were almost unidentifiable whereas the performance on others was above chance level.

#### 3.1.2. Experiment 2

The overall performance was 46% for the original MRIs, 25% for PyDeface, 29% for AnonyMI and 30% for Maskface. All three deidentification methods reduced the probability of correct responses in a statistically significant manner. PyDeface provided the highest reduction in performance with respect to the original MRIs (β = -0.92, *p* < .001), followed by AnonyMI (β = - 0.70, *p* < .001) and Maskface (β = -0.67, *p* < .001). See Supplementary Table 2 for a complete report of this analysis. The overall performance across stimulus subjects varied from 15% to 46% (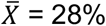, *sd* = 8%), showing again that there were subjects that were almost unidentifiable whereas others were above chance level.

### 3.2. MRI Geometrical Similarity and Application to Source Localization

#### 3.2.1. Geometrical Similarity

The Jaccard Distance on skull-stripped images analysis showed that AnonyMI resulted in a mean similarity of 0.9982 and for Maskface of 0.9985 with respect to the original MRIs, whereas PyDeface resulted in 0.9686 (with some volumes going below 0.9). The differences between methods were all statistically significant (p < 0.001), however, the difference of 0.0001 between AnonyMI and Maskface holds little practical significance, especially if compared to the difference of ∼0.03 between both of these methods and PyDeface.

The analysis on the 3D reconstructions of the brain, skull and skin surfaces revealed that the original MRIs, AnonyMI and PyDeface yielded 92% of usable models (all three failing on the exact same MRIs), whereas Maskface produced 69%. This difference was statistically significant as assessed by McNemar’s Chi-squared test (χ^2^= 13.06; *p* < .001).

The pairwise analysis of the Hausdorff Distance by surface and method showed significant differences between PyDeface (*<u>X</u>* = 49.3) vs. AnonyMI (*<u>X</u>* = 4.69; *p* < .001) and PyDeface vs. Maskface (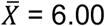; *p* < .001) for the outer skull surface; and between PyDeface 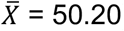 vs. AnonyMI (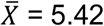; *p* < .001), PyDeface vs. Maskface (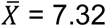; *p* < .001) and AnonyMI vs. Maskface (*p* < .05) for the outer skin surface. See Supplementary Table 3 for a complete report of this analysis. Overall, these results showed that AnonyMI provided the highest level of geometrical preservation.

#### 3.2.1. Source Localization

The source localization analysis showed that AnonyMi provided 88% of equal solutions with respect to the results obtained with the original MRIs, while Maskface provided 85% and PyDeface 84% (Figure 3.f). The difference between methods was statistically significant as assessed by McNemar’s Chi-squared tests. The overall distance between the real sources and the reconstructed ones was not statistically between methods, due to the relatively high number of equal solutions yielded by each method (> 84%). However, all of them differed significantly from the results obtained with the original MRIs (Figure 3.g). The overall distance between the sources reconstructed with the original MRI and the sources reconstructed with each method is shown in Figure 3.h for each solution in which at least one method showed a different result (same number of observations across methods), and on Figure 3.i for the solutions on which each of the methods showed a different results (different number of observations across methods).

## 4. Discussion

Deidentifying MRIs constitutes a challenge as it has to balance two opposite goals. It should prevent the possibility of re-identification of a participant while keeping as much geometrical information as possible. To tackle this issue, we developed a novel MRI deidentification method that aims at providing a balance between these objectives. In order to assess the performance on these two fronts, and to compare it to other state-of-the-art methods we performed two series of analysis: a behavioral and a geometrical validation.

To the best of our knowledge, this is the first study that systematically evaluated and compared the performance of state-of-the-art MRI deidentification methods. In two behavioral experiments we tested how well could participants identify subjects from 3D reconstructions of their MRIs. On Experiment 1, where subjects were first presented with a single photograph which was followed by four deidentified MRIs, the performance was 33, 37 and 40% for PyDeface, AnonyMI and Maskface, respectively. The only statistically significant difference was found between PyDeface and Maskface. Even though the average results were above chance level (25%), the general performance of all three methods can be arguably considered satisfactory, as the experimental design corresponded more to a discrimination task than to an identification task, which would have been more difficult. In other words, if subjects were able to discard one of the four testing images and guess randomly, chance-level would rise to ∼33%, which is close to the observed performance levels. To complement these results, we performed a second experiment, with a more ecological design, in which participants were presented with four photographs and then with a single reconstructed MRI and had to decide to which of the four photographs it corresponded. This task included a memory component as subjects had to remember the faces and reduced the possibility of using a discrimination strategy, which made it more similar to a real-life situation. In this case, the performance was 25, 29 and 30%, for PyDeface, AnonyMI and Maskface, respectively. In this task we also included a control block on which the MRIs were not deidentified, which allowed us to estimate a baseline level of identification performance, which was 46%. It is worth noting that the performance with the original MRIs might be inflated with respect to the anonymized ones as this block was always carried out at the end of the task, when subjects had had more training. This experimental design was chosen in order to avoid that subjects found distinctive features on the non-deidentified stimuli that could be used on deidentified trials and bias the results of the main aim of the experiment. The differences among methods were not statistically significant, but the identification performance of all three methods was significantly lower than in the non-deidentified condition. Interestingly, in both behavioral experiments the analysis on individual stimulus subjects showed that different methods worked better with different subjects. This overall pattern of results indicates that even though PyDeface was the most performant deidentification method in terms of reducing the re-identification risk, followed by AnonyMi and finally by Maskface, all three methods efficiently reduced the risk of identification almost at chance level. This notion is of paramount importance in light of the new legislation for data privacy. Indeed, if it is assumed that all currently used methods are sufficient to appropriately deidentify MRI data, that is, that it is not possible to re-identify a data subject from such a “deidentified” dataset, and this assumption is incorrect then it leaves open the possibility of a major data breach. Data breaches can seriously infringe the privacy of patients and research subjects, and also, under GDPR, they can have significant financial implications for the offending scientist and their institution. Administrative fines are detailed in Article 83 of the GDPR and stipulate that, depending upon the severity of the breach, fines could be up to €20 million or 4% of the institutions’ global annual turnover of the previous financial year, whichever is higher (European Parliament and Council of European Union, 2016). This is not to mention the potential reputational damage and distress to data subjects such a breach might cause.

Our study focused on the identification risk by a human operator, however, a growing body of literature shows that machine learning algorithms could potentially be used to individuate research participants from shared data and even to reconstruct the subject’s face from deidentified MRIs (Abramian and Eklund, 2019; Schwarz et al., 2019). An in-depth analysis of this risk is of utmost importance for future studies in order to provide a comprehensive picture of the risks associated with making deidentified MRIs public.

While, in the case of data sharing, reducing the re-identification risk of the MRI is obviously the main aim of any MRI deidentification method, the associate distortions of the geometrical feature of the images should be maximally reduced in order to maximize reuse and interoperability (Wilkinson et al., 2016). For this reason, we performed a spatial analysis in which we tested how well each method preserved the geometrical characteristics of the MRIs. We first tested the Jaccard Similarity of skull stripped brain volumes done with the deidentified MRIs with respect those obtained with the original MRIs. In this analysis Maskface and AnonyMI showed ∼99% convergence, whereas PyDeface showed 96%, with some images going below 90% with the latter. It is worth mentioning that the skull stripping procedure was carried out with a defaced template in the case of the PyDeface method while it was done with a full-face template for Maskface and AnonyMI, because using the latter for PyDeface would have unfairly underestimated its performance. These results indicate that both Maskface and AnonyMI provide deidentified volumes that give more similar results to those obtained with the original MRIs when applying a subsequent processing algorithm as skull stripping. This is important to ensure replicability of results when using deidentified images.

Next, we extracted 3D surfaces of the brain, skull and skin using the watershed algorithm, a standard procedure for performing M/EEG source localization. We found that Maskface provided 30% of unusable surfaces due to severe geometrical distortions, whereas with PyDeface and AnonyMI it was 8%, the same percentage obtained with the original MRIs. Then, employing the successfully reconstructed surfaces we computed how similar they were to the surfaces obtained with the original MRIs. This analysis showed that AnonyMI significantly outperformed PyDeface for the outer skull, and also outperformed both PyDeface and Maskface for the skin. This pattern of results indicates that AnonyMI was the method that induced less geometrical distortions.

To assess how much using deidentified MRIs affects source localization results, we reconstructed electrical sources using a dataset with ground-truth and compared the results obtained with each deidentification method to those obtained with the original MRIs. This analysis showed that the percentage of equal solution with respect to the original MRIs was 88, 85 and 84% for AnonyMI, Maskface and PyDeface, respectively. In other words, AnonyMI provided an increase of ∼4% of equal solution across multiple parameter configurations. The overall pattern of geometrical results shows that AnonyMI provides the most reliable MRIs for performing structural and source localization analysis. The increase in the number of equal solutions might be more pronounced if using FEM instead of BEM models (Hallez et al., 2007), as the former creates a more detailed geometrical reconstruction. We chose to use the BEM method, as it is arguably the most widely used method for distributed source modelling due to its balance of low computational costs with geometrical correctness (Hallez et al., 2007).

Aside from the aforementioned benefits, AnonyMI presents many other advantages. It leverages the power of 3D Slicer for importing and saving in any filetype, avoiding unnecessary format changes that are mandatory with other methods and that can lead to errors. It also takes advantage of 3D Slicer for fast and practical visualization of results, which eases the quality assurance of the procedure. It can also create population specific templates in minutes, simplifying the analysis of particular datasets (i.e. age specific). It provides a visual interface to mask specific areas of a subject that might have severely increased the re-identification risk. It can easily run multiple subjects automatically, making it easy to process large datasets and has both a command-line and a graphical interface, providing an essential solution for non-programmers.

In sum, AnonyMI is a novel method that offers several technical advantages that make MRI deidentification easier, and that provides the optimal balance between deidentification performance and geometrical preservation.

## 5. Limitations

The present study is the first to comparatively analyze the performance of different MRI deidentification methods, however, further studies will be required to obtain a more comprehensive characterization. We focused on two particular behavioral tasks to assess the identification risk and to quantify the improvements provided by each method, aiming at tasks that could provide ecologically valid estimates. Nevertheless, different tasks might provide different results, and therefore, more experiments with different experimental designs will be required to thoroughly describe each method’s performance. Similarly, the stimuli employed, and the participants of the behavioral tasks corresponded to relatively homogeneous samples of white caucasian subjects, therefore the generalization of our results to other ethnicities is not granted and should be addressed in future studies (Dotson and Duarte, 2020). With regards to the geometrical analysis, we focused on three measures (Jaccard similarity, Hausdorff distance and BEM outcome). A comparison of more measures and analysis pipelines would be of great importance to appraise how much each method allows reusability under different circumstances. Finally, our source localization analysis focused on one forward model type (i.e. BEM), three inverse solution methods (dSPM, eLORETA and MNE), and used non-physiological ground truth data. We chose these methods as they are among the most used methods in contemporary M/EEG source localization analysis. However, an investigation of other forward models and inverse solution methods would provide a more nuanced comparison.

## 6. Author Contributions

**Ezequiel Mikulan**: Conceptualization, Methodology, Software, Formal analysis, Writing - Original Draft. **Simone Russo**: Investigation, Validation. **William Knight**: Writing - Original Draft. **Flavia Maria Zauli**: Investigation, Data Curation. **Sara Parimigiani**: Investigation, Validation. **Jacopo Favaro**: Investigation, Validation **d’Orio Piergiorgio**: Investigation, Validation. **Silvia Squarza**: Validation. **Pierluigi Perri**: Writing - Original Draft. **Francesco Cardinale**: Investigation, Validation, Writing - Review & Editing. **Pietro Avanzini**: Conceptualization, Methodology, Writing - Original Draft, Writing - Review & Editing. **Andrea Pigorini**: Conceptualization, Methodology, Writing - Original Draft, Writing - Review & Editing, Project administration, Supervision.

## References

Abramian, D., Eklund, A., 2019. Refacing: Reconstructing Anonymized Facial Features Using GANS, in: 2019 IEEE 16th International Symposium on Biomedical Imaging (ISBI 2019). Presented at the 2019 IEEE 16th International Symposium on Biomedical Imaging (ISBI 2019), pp. 1104–1108. https://doi.org/10.1109/ISBI.2019.8759515

Akalin-Acar, Z., Gençer, N.G., 2004. An advanced boundary element method (BEM) implementation for the forward problem of electromagnetic source imaging. Phys. Med. Biol. 49, 5011–5028. https://doi.org/10.1088/0031-9155/49/21/012

Amunts, K., Knoll, A.C., Lippert, T., Pennartz, C.M.A., Ryvlin, P., Destexhe, A., Jirsa, V.K., D’Angelo, E., Bjaalie, J.G., 2019. The Human Brain Project—Synergy between neuroscience, computing, informatics, and brain-inspired technologies. PLOS Biol. 17, e3000344. https://doi.org/10.1371/journal.pbio.3000344

Ascoli, G.A., Maraver, P., Nanda, S., Polavaram, S., Armañanzas, R., 2017. Win–win data sharing in neuroscience. Nat. Methods 14, 112–116. https://doi.org/10.1038/nmeth.4152

Avants, B., Tustison, N., 2018. ANTs/ANTsR Brain Templates. https://doi.org/10.6084/m9.figshare.915436.v2

Avants, B.B., Tustison, N.J., Song, G., Cook, P.A., Klein, A., Gee, J.C., 2011. A reproducible evaluation of ANTs similarity metric performance in brain image registration. NeuroImage 54, 2033–2044. https://doi.org/10.1016/j.neuroimage.2010.09.025

Bates, D., Mächler, M., Bolker, B., Walker, S., 2015. Fitting Linear Mixed-Effects Models Using lme4. J. Stat. Softw. 67, 1–48. https://doi.org/10.18637/jss.v067.i01

Bischoff-Grethe, A., Ozyurt, I.B., Busa, E., Quinn, B.T., Fennema-Notestine, C., Clark, C.P., Morris, S., Bondi, M.W., Jernigan, T.L., Dale, A.M., Brown, G.G., Fischl, B., 2007. A Technique for the Deidentification of Structural Brain MR Images. Hum. Brain Mapp. 28, 892–903. https://doi.org/10.1002/hbm.20312

Brakewood, B., Poldrack, R.A., 2013. The ethics of secondary data analysis: Considering the application of Belmont principles to the sharing of neuroimaging data. NeuroImage 82, 671–676. https://doi.org/10.1016/j.neuroimage.2013.02.040

Budin, F., Zeng, D., Ghosh, A., Bullitt, E., 2008. Preventing facial recognition when rendering MR images of the head in three dimensions. Med. Image Anal. 12, 229–239. https://doi.org/10.1016/j.media.2007.10.008

Dale, A.M., Fischl, B., Sereno, M.I., 1999. Cortical Surface-Based Analysis: I. Segmentation and Surface Reconstruction. NeuroImage 9, 179–194. https://doi.org/10.1006/nimg.1998.0395

de Sitter, A., Visser, M., Brouwer, I., Cover, K.S., van Schijndel, R.A., Eijgelaar, R.S., Müller, D.M.J., Ropele, S., Kappos, L., Rovira, Á., Filippi, M., Enzinger, C., Frederiksen, J., Ciccarelli, O., Guttmann, C.R.G., Wattjes, M.P., Witte, M.G., de Witt Hamer, P.C., Barkhof, F., Vrenken, H., on behalf of the MAGNIMS Study Group and Alzheimer’s Disease Neuroimaging Initiative, 2020. Facing privacy in neuroimaging: removing facial features degrades performance of image analysis methods. Eur. Radiol. 30, 1062–1074. https://doi.org/10.1007/s00330-019-06459-3

Dixon, P., 2008. Models of accuracy in repeated-measures designs. J. Mem. Lang. 59, 447– 456. https://doi.org/10.1016/j.jml.2007.11.004

Dotson, V.M., Duarte, A., 2020. The importance of diversity in cognitive neuroscience. Ann. N. Y. Acad. Sci. 1464, 181–191. https://doi.org/10.1111/nyas.14268

European Parliament, Council of European Union, 2016. Regulation (EU) 2016/679.

Fedorov, A., Beichel, R., Kalpathy-Cramer, J., Finet, J., Fillion-Robin, J.-C., Pujol, S., Bauer, C., Jennings, D., Fennessy, F., Sonka, M., Buatti, J., Aylward, S., Miller, J.V., Pieper, S., Kikinis, R., 2012. 3D Slicer as an image computing platform for the Quantitative Imaging Network. Magn. Reson. Imaging, Quantitative Imaging in Cancer 30, 1323–1341. https://doi.org/10.1016/j.mri.2012.05.001

Gulban, O.F., Nielson, D., Poldrack, R., Lee, J., Gorgolewski, C., Vanessasaurus Ghosh, S., 2019. poldracklab/pydeface: v2.0.0. Zenodo. https://doi.org/10.5281/zenodo.3524401

Hallez, H., Vanrumste, B., Grech, R., Muscat, J., De Clercq, W., Vergult, A., D’Asseler, Y., Camilleri, K.P., Fabri, S.G., Van Huffel, S., Lemahieu, I., 2007. Review on solving the forward problem in EEG source analysis. J. NeuroEngineering Rehabil. 4, 46. https://doi.org/10.1186/1743-0003-4-46

Hamalainen, M.S., Sarvas, J., 1987. Feasibility of the homogeneous head model in the interpretation of neuromagnetic fields. Phys. Med. Biol. 32, 91–97. https://doi.org/10.1088/0031-9155/32/1/014

Huttenlocher, D.P., Klanderman, G.A., Rucklidge, W.J., 1993. Comparing images using the Hausdorff distance. IEEE Trans. Pattern Anal. Mach. Intell. 15, 850–863. https://doi.org/10.1109/34.232073

Jaccard, P., 1912. The Distribution of the Flora in the Alpine Zone.1. New Phytol. 11, 37–50. https://doi.org/10.1111/j.1469-8137.1912.tb05611.x

Jaeger, T.F., 2008. Categorical Data Analysis: Away from ANOVAs (transformation or not) and towards Logit Mixed Models. J. Mem. Lang. 59, 434–446. https://doi.org/10.1016/j.jml.2007.11.007

Kalavathi, P., Prasath, V.B.S., 2016. Methods on Skull Stripping of MRI Head Scan Images—a Review. J. Digit. Imaging 29, 365–379. https://doi.org/10.1007/s10278-015-9847-8

Kalkman, S., Mostert, M., Gerlinger, C., van Delden, J.J.M., van Thiel, G.J.M.W., 2019. Responsible data sharing in international health research: a systematic review of principles and norms. BMC Med. Ethics 20, 21. https://doi.org/10.1186/s12910-019-0359-9

Kushida, C.A., Nichols, D.A., Jadrnicek, R., Miller, R., Walsh, J.K., Griffin, K., 2012. Strategies for de-identification and anonymization of electronic health record data for use in multicenter research studies. Med. Care 50, S82–101. https://doi.org/10.1097/MLR.0b013e3182585355

Mazura, J.C., Juluru, K., Chen, J.J., Morgan, T.A., John, M., Siegel, E.L., 2012. Facial Recognition Software Success Rates for the Identification of 3D Surface Reconstructed Facial Images: Implications for Patient Privacy and Security. J. Digit. Imaging 25, 347– 351. https://doi.org/10.1007/s10278-011-9429-3

Mikulan, E., Russo, S., Parmigiani, S., Sarasso, S., Zauli, F.M., Rubino, A., Avanzini, P., Cattani, A., Sorrentino, A., Gibbs, S., Cardinale, F., Sartori, I., Nobili, L., Massimini, M., Pigorini, A., 2020. Simultaneous human intracerebral stimulation and HD-EEG, ground-truth for source localization methods. Sci. Data 7, 1–8. https://doi.org/10.1038/s41597-020-0467-x

Milchenko, M., Marcus, D., 2013. Obscuring Surface Anatomy in Volumetric Imaging Data. Neuroinformatics 11, 65–75. https://doi.org/10.1007/s12021-012-9160-3

Milham, M.P., Craddock, R.C., Son, J.J., Fleischmann, M., Clucas, J., Xu, H., Koo, B., Krishnakumar, A., Biswal, B.B., Castellanos, F.X., Colcombe, S., Di Martino, A., Zuo, X.-N., Klein, A., 2018. Assessment of the impact of shared brain imaging data on the scientific literature. Nat. Commun. 9, 2818. https://doi.org/10.1038/s41467-018-04976-1

Mott, M.C., Gordon, J.A., Koroshetz, W.J., 2018. The NIH BRAIN Initiative: Advancing neurotechnologies, integrating disciplines. PLOS Biol. 16, e3000066. https://doi.org/10.1371/journal.pbio.3000066

Peirce, J., Gray, J.R., Simpson, S., MacAskill, M., Höchenberger, R., Sogo, H., Kastman, E., Lindeløv, J.K., 2019. PsychoPy2: Experiments in behavior made easy. Behav. Res. Methods 51, 195–203. https://doi.org/10.3758/s13428-018-01193-y

Poline, J.-B., Breeze, J.L., Ghosh, S., Gorgolewski, K., Halchenko, Y.O., Hanke, M., Haselgrove, C., Helmer, K.G., Keator, D.B., Marcus, D.S., Poldrack, R.A., Schwartz, Y., Ashburner, J., Kennedy, D.N., 2012. Data sharing in neuroimaging research. Front. Neuroinformatics 6, 9. https://doi.org/10.3389/fninf.2012.00009

Prior, F.W., Brunsden, B., Hildebolt, C., Nolan, T.S., Pringle, M., Vaishnavi, S.N., Larson-Prior, L.J., 2009. Facial Recognition From Volume-Rendered Magnetic Resonance Imaging Data. IEEE Trans. Inf. Technol. Biomed. 13, 5–9. https://doi.org/10.1109/TITB.2008.2003335

R Core Team, 2019. R: A Language and Environment for Statistical Computing. R Foundation for Statistical Computing, Vienna, Austria.

Schwarz, C.G., Kremers, W.K., Therneau, T.M., Sharp, R.R., Gunter, J.L., Vemuri, P., Arani, A., Spychalla, A.J., Kantarci, K., Knopman, D.S., Petersen, R.C., Jack, C.R., 2019. Identification of Anonymous MRI Research Participants with Face-Recognition Software. N. Engl. J. Med. 381, 1684–1686. https://doi.org/10.1056/NEJMc1908881

Ségonne, F., Dale, A.M., Busa, E., Glessner, M., Salat, D., Hahn, H.K., Fischl, B., 2004. A hybrid approach to the skull stripping problem in MRI. NeuroImage 22, 1060–1075. https://doi.org/10.1016/j.neuroimage.2004.03.032

Souza, R., Lucena, O., Garrafa, J., Gobbi, D., Saluzzi, M., Appenzeller, S., Rittner, L., Frayne, R., Lotufo, R., 2018. An open, multi-vendor, multi-field-strength brain MR dataset and analysis of publicly available skull stripping methods agreement. NeuroImage, Segmenting the Brain 170, 482–494. https://doi.org/10.1016/j.neuroimage.2017.08.021

Tustison, N.J., Cook, P.A., Klein, A., Song, G., Das, S.R., Duda, J.T., Kandel, B.M., van Strien, N., Stone, J.R., Gee, J.C., Avants, B.B., 2014. Large-scale evaluation of ANTs and FreeSurfer cortical thickness measurements. NeuroImage 99, 166–179. https://doi.org/10.1016/j.neuroimage.2014.05.044

Wilkinson, M.D., Dumontier, M., Aalbersberg, Ij.J., Appleton, G., Axton, M., Baak, A., Blomberg, N., Boiten, J.-W., da Silva Santos, L.B., Bourne, P.E., Bouwman, J., Brookes, A.J., Clark, T., Crosas, M., Dillo, I., Dumon, O., Edmunds, S., Evelo, C.T., Finkers, R., Gonzalez-Beltran, A., Gray, A.J.G., Groth, P., Goble, C., Grethe, J.S., Heringa, J., ‘t Hoen, P.A.C., Hooft, R., Kuhn, T., Kok, R., Kok, J., Lusher, S.J., Martone, M.E., Mons, A., Packer, A.L., Persson, B., Rocca-Serra, P., Roos, M., van Schaik, R., Sansone, S.-A., Schultes, E., Sengstag, T., Slater, T., Strawn, G., Swertz, M.A., Thompson, M., van der Lei, J., van Mulligen, E., Velterop, J., Waagmeester, A., Wittenburg, P., Wolstencroft, K., Zhao, J., Mons, B., 2016. The FAIR Guiding Principles for scientific data management and stewardship. Sci. Data 3, 160018. https://doi.org/10.1038/sdata.2016.18

